# Contrastive Learning for Robust Cell Annotation and Representation from Single-Cell Transcriptomics

**DOI:** 10.1101/2024.06.20.599868

**Authors:** Leo Andrekson, Rocío Mercado

## Abstract

Batch effects are a significant concern in single-cell RNA sequencing (scRNA-Seq) data analysis, where variations in the data can be attributed to factors unrelated to cell types. This can make downstream analysis a challenging task. In this study, we present a novel DL approach using contrastive learning and a carefully designed loss function for learning a generalizable embedding space from scRNA-Seq data. We call this model CELLULAR: CELLUlar contrastive Learning for Annotation and Representation. When benchmarked against multiple established methods for scRNA-Seq integration, CELLULAR outperforms existing methods in learning a generalizable embedding space on multiple datasets. Cell annotation was also explored as a downstream application for the learned embedding space. When compared against multiple well-established methods, CELLULAR demonstrates competitive performance with top cell classification methods in terms of accuracy, balanced accuracy, and F1 score. CELLULAR is also capable of performing novel cell type detection. These findings assess the biological relevance of the model’s learned embedding space by demonstrating the robustness of its cell representations across various applications. The model has been structured into an opensource Python package, specifically designed to simplify and streamline its usage for bioinformaticians and other scientists interested in cell representation learning.

## 1 Introduction

Cells in our body exhibit remarkable diversity, and characterizing and classifying different cell types is fundamental to understanding their functions and roles in various biological processes.(1) Learning compact vector representations of cell types enables us to algorithmically organize and categorize cells based on shared characteristics, such as their observed phenotypes, within a lower-dimensional space than that afforded by measurements of cellular phenotypes. Also known as *cell representation learning*, this has broad implications in biomedical research, including disease diagnostics and drug development, and has become an active research area intersecting deep learning (DL) and biology. Such compact representations would enable us to uncover previously unknown relationships between cells, identify disease-specific cell populations, and design targeted therapies for specific cell types and tissues; nevertheless, no generalizable approach exists which can effectively achieve this currently.

Towards the aim of learning compact and generalizable representations of cells, this study leverages singlecell RNA sequencing (scRNA-Seq) data to train a neural network to produce an efficient embedding space for unique cell types. The method is called CELLULAR, or CELLUlar contrastive Learning for Annotation and Representation, and presented as an open-source Python package. We benchmark CELLULAR against diverse leading methods for scRNA-Seq data integration, and demonstrate the utility of the learned embedding space on two downstream applications: cell type annotation and novel cell type detection. This enabled us to assess CELLULAR’s performance by validating its latent space for key bioinformatics tasks in single-cell transcriptomics.

## 2 Background

### 2.1 Single-cell omics

In the last decade, single-cell omics (sc-omics) has emerged as a rapidly growing field of biological research due to its applications in atlasing projects (e.g., the Human Cell Atlas), developmental studies, and precision medicine. Popular single-cell methods include transcriptomics, epigenomics, and proteomics, where scRNA-Seq (Figure 1a), a transcriptomics approach, is currently the most widely used sc-omics method.(1, 2) With the rise of single-cell technologies, multi-omics analysis has gained prominence, enabling the integration of diverse omics data to uncover complex cellular properties.(3) Common approaches include transcriptomics-genomics and transcriptomics-epigenomics integration.(4) As these technologies advance and become more accessible, they offer critical insights into disease progression and treatment responses at single-cell resolution, key for personalized medicine.(1) However, effectively leveraging sc-omics data requires overcoming challenges in high noise and dimensionality, underscoring the need for DL models that learn a generalizable latent space from vast amounts of noisy, single-cell data.

**Fig. 1.**
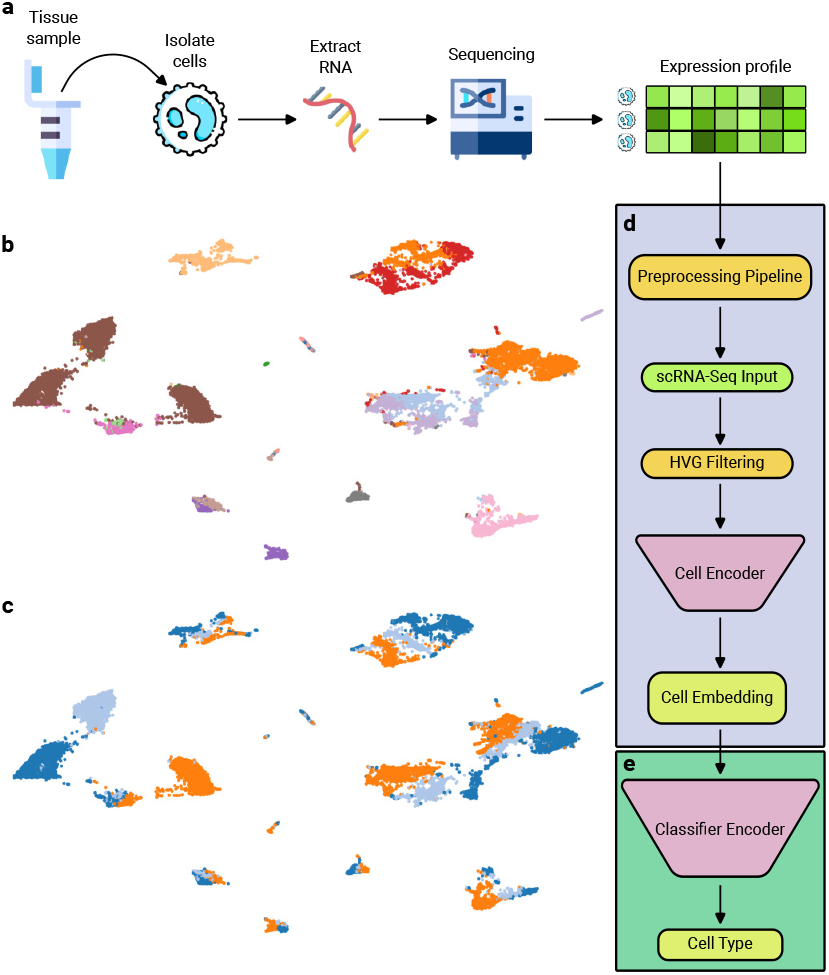
(a) ScRNA sequencing generally consists of sampling, isolation of cells, RNA extraction, reverse transcription to cDNA, sequencing, and eventual data analysis. (b) UMAP visualization of the kidney scRNA-Seq dataset constructed in this study, colored by cell type. (b) UMAP visualization colored by patient ID, highlighting batch effects due to patient ID. (d) Overview of CELLULAR: ScRNA-Seq data are fed through a feed-forward encoder that returns a learned embedding for each sample (cell). (e) CELLULAR embeddings can then be used as input to another feed-forward neural network to classify samples into potential cell types.

#### 2.1.1 Cell type annotation

As large-scale sc-omics become more accessible, demand for automated cell annotation has increased due to the limitations of manual annotation, which is time consuming and subjective. Recently developed strategies for the annotation of cell types from scRNA-Seq data typically rely on referencing marker gene databases (5, 6), correlating reference expression data (7, 8), or supervised classification methods.(9–11) Here, we present a novel method for supervised cell type annotation. The model learns an embedding space from labeled scRNA-Seq data through contrastive learning (CL) at training time, and subsequently uses this space for cell type classification at inference.

#### 2.1.2 ScRNA-Seq data integration

ScRNA-Seq datasets often encompass numerous samples, leading to the inevitable introduction of batch effects arising from, e.g., diverse patients or sequencing methodology.(12) Here, batch effects denote sources of variation within the data attributed to non-biological factors. To emphasize the challenges imposed by batch effects, we visualize some example data in Figure 1b&c. The ideal model would integrate scRNA-Seq data into an embedding space where biological variation is maximized while minimizing variation from batch effects— easier said than done. Many methods have been developed for this purpose, such as scANVI (13) and scGen (9), which have been benchmarked in a recent study (14). While these methods show promise in scRNA-Seq data integration, they were not evaluated for generalization of the embedding space.

The scRNA-Seq integration benchmark introduced by Luecken et al. (14), aptly named *single-cell integration benchmarking* or *scIB*, also compared approaches utilizing transformers (TOSICA (11)), variational autoencoders (VAEs; scGen (9)), and hierarchical Bayesian models (scVI (15), scANVI (13)). By comparing these methods with CELLULAR, we can evaluate how CL fares on data integration benchmarks. By including multiple modeling approaches that span the breadth of previous research in this area, we ensure a fair and comprehensive comparison of our method to prior work. Many other approaches exist, such as Scanorama (16) and Harmony (17), but are not able to make predictions on novel data; most models have been developed to only integrate existing data.

In Figure 1c we see how variation between patients leads to separate clusters forming for data from cells of the same type (Figure 1b). This is not desirable and patient ID would be considered a batch effect in this scenario; instead, cells of the same type should cluster together such that the data become completely mixed in terms of patient ID, and no variation within the dataset should be attributable to batch effects (e.g., patient, laboratory). The data in Figure 1 all come from healthy individuals, so the patient variable would ideally not capture significant biological variation. Nevertheless, attaining complete mitigation of batch effects is a significant challenge in scRNA-Seq data integration.

### 2.2 Previous work

#### 2.2.1 ScRNA-Seq integration

There has been significant previous work for scRNA-Seq integration. Here, we compare CELLULAR to the following studies: TOSICA (11), scGen (9), scANVI (13), and scVI (15). The models we chose to compare here were selected for their good generalization capabilities previously demonstrated on new data and diverse architectures, allowing us to compare a broad range of leading methods to further evaluate their generalizability.

CELLULAR is most similar to TOSICA and scGen. TOSICA uses transformers for cell type annotation from scRNA-Seq data; it takes advantage of pathway and regulon information for learning. scGen uses VAEs to predict cellular responses to perturbations from scRNA-Seq data. On the other hand, scVI and scANVI both use hierarchical Bayesian models where conditional distributions are learned via neural networks; these models have the advantage of efficient training, even for very large datasets. As we demonstrate in this study, although CELLULAR uses a simpler neural network architecture than TOSICA, scVI, and scANVI, it achieves superior performance to these models on generalizable data integration, and modestly better performance to scGen. Nevertheless, the performance of all of these models on data integration is very dataset-dependent. We do not explore the effect of perturbations on cellular phenotype in this work.

#### 2.2.2 Cell type annotation

There has also been notable previous work for cell type annotation from scRNA-Seq data. In this work, we compare CELLULAR to the following studies: TOSICA (11), scNym (18), Seurat (19), SciBet (20), and CellID (21). While there are many published models for cell type annotation, we specifically selected these five because they cover a wide range of approaches for solving cell annotation tasks. For instance, TOSICA is a supervised algorithm, principle component analysis (PCA) is an unsupervised dimensionality reduction method, and scANVI is a semi-supervised approach. scNym is an adversarial neural network trained in a semi-supervised way for cell type classification from a variety of sc-omics data, and can also be used for novel cell type detection. Seurat combines dictionary learning and dataset sketching for the analysis of large scale sc-transcriptomics data. SciBet uses multinomial logistic regression for cell annotation from scRNA-Seq data. Finally, CellID is a clustering-free multivariate statistical approach to cell annotation from sc-omics data. In this setting, CELLULAR is most similar to TOSICA and scNym.

In the annotation benchmark presented here, we evaluate the aforementioned models in two ways: first, by training the models using all gene information (in models that can handle this); and second, by filtering the gene set to use only highly variable genes (HVGs) in training. In prior work, only TOSICA used HVG filtering, but by doing HVG filtering on all models, we can enable a more fair comparison.

Other related work to which we do not compare to in this work includes Richter et al. (22), where the authors used self-supervised learning techniques for data integration and cell classification, amongst other tasks. In that study, concurrent to this one, the authors also found that integration of CL improved the generalizability of their method on sc-genomics data. There are also scPoli (23) and scGPT (24), two methods which handle both integration and classification, CellHint (25), a clustering tree-based model for cell type annotation, and Symphony (26). CAMLU (27) is an R package that uses autoencoders for novel cell type detection.

### 2.3 Motivation

Many dimensionality reduction methods exist, broadly categorized into statistical methods, manifold methods, and neural networks.(28) This study focuses on developing a neural network approach to generate generalizable cell type representations while minimizing batch effects and preserving biological variation. Prior neural network-based batch correction methods have used autoencoders and VAEs (9, 22, 23, 27, 29, 30), generative adversarial networks (GANs) (31), recurrent neural networks (RNNs) (32), contrastive learning (CL) (22, 33, 34), and transformers (11, 24).

CELLULAR introduces a novel combination of CL (Equation 7) and a cell type centroid loss (Equation 8), specifically designed for sc-omics integration. The contrastive loss encourages embeddings of the same cell type to cluster together, while the centroid loss maintains relative positioning between clusters. Cell type centroids are computed using PCA, chosen for its linearity, interpretability, and ability to preserve variance, unlike non-linear methods like UMAP which can distort global relationships. We demonstrate that this approach enhances robustness to noise and batch effects, and produces biologically meaningful embeddings.

## 3 Methods

### 3.1 Model architecture

CELLULAR consists of a feed-forward encoder that can be trained on scRNA-Seq data to produce a latent space (Figure 1d). The encoder consists of 2 linear layers, each followed by layer normalization and ReLU activation. The encoder is designed to compress the input after each layer, ending with a final embedding space of dimension 100, which can be used as input for any downstream neural network, such as a classifier (Figure 1e). CELLULAR contains 2,558,600 learnable weights under default settings, and uses the Adam optimizer for training with a cosine warm-up scheduler. CELLULAR was trained on an NVIDIA A100 GPU using randomly-sampled mini batches of size 256. For both of the downstream tasks explored herein—cell annotation and novel cell type classification—the initial encoder weights are fixed. Only the classifier model weights are updated when training the model for cell type annotation.

### 3.2 Data

Data collected from human participants were gathered from the following studies and tissues:

- Bone marrow: GSM3396161, GSM3396176, GSM3396184, GSM3396183 and GSM3396174.(35) Data available for 20 cell types and 5 patients.
- PBMC: GSE115189, GSM3665016, GSM3665017, GSM3665018, GSM3665019 and PBMC 10X data.(36, 37) Data available for 17 cell types and 6 patients.
- Pancreas: GSE81076, GSE85241, GSE86469 and E-MTAB-5061, gathered into one dataset by the Satija lab (38). Data available for 9 cell types and 4 sources. It is unclear if data comes only from different sequencing methods or also different patients; for simplicity, we label the source *patient ID*.
- Kidney: GSM4572192, GSM4572193 and GSM4572194.(39) Data available for 15 cell types and 3 patients.
- Merged: Bone marrow, PBMC, pancreas, and kidney datasets all merged into one dataset consisting of 46 cell types and 18 patients.

These datasets were used to assess the models’ capabilities in creating a generalizable embedding spaces from scRNA-Seq data. For exploring the models’ cell type annotation capabilities, the following datasets were used:

- Baron: GSE84133.(40) Data available for 14 cell types and 4 patients.
- MacParland: GSE115469.(41) Data available for 20 cell types and 5 patients.
- Segerstolpe: E-MTAB-5061.(42) Data available for 15 cell types and 10 patients.
- Zheng68k: Data from the ‘Fresh 68K PBMCs’ section at 10xGenomics.(43) Data available for 11 cell types and 8 unique barcode identifiers.

After preprocessing, each of the aforementioned datasets contains the following number of cells: bone marrow—14,779; PBMC—22,027; kidney—16,370; pancreas—6,313; merged dataset—59,489; Baron— 8,569; MacParland—8,444; Segerstolpe—3,514; and Zheng68k—68,579.

### 3.3 Preprocessing

ScRNA-Seq data were preprocessed as follows:

1. Cells filtered based on the number of unique genes required for downstream processing by applying a shifted logarithm transformation and removing cells that exceeded a threshold of 5 median absolute deviations (MAD) from the median, where *MAD* = *median*(|*x* − *median*(*x*)|).
2. Cells filtered by sequencing depth to ensure highquality samples using the shifted logarithm of read counts. Samples were again filtered using a MAD threshold of 5.
3. Cells filtered by the proportion of mitochondrial counts. This was done to remove cells suspected of having broken membranes, which could potentially be dying cells.(44) Filtering was performed using a MAD threshold of 5 for almost all datasets, occasionally changed to a threshold between 5 and 10 based on a visual assessment of the distributions.
4. Cells filtered by the proportion of total counts corresponding to the top 20 genes with the highest number of counts per cell. Filtering was performed using a MAD threshold of 5 for almost all datasets, occasionally changed to a threshold between 3 and 5 based on a visual assessment.
5. Genes filtered by number of unique cells expressing a gene. Genes that appeared in less than 20 cells were discarded.
6. Normalization was done in 2 steps. First, total counts were normalized to ensure comparability across cells by computing the average count per cell,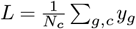, where *N*_*c*_ is the number of cells and *y* is the counts matrix; index *g* annotates the position of a gene and *c* the position of a cell. Each cell’s total count was then scaled by *L*, yielding 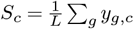. Next, to approximate a normal distribution, a shifted logarithm transformation—shown to perform well in recent benchmarks (45)—was applied:

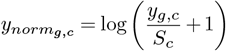

where *y*_*norm*_ is the normalized counts matrix.
7. Cell type annotation was performed using *ScType*.(6)
8. Finally, we select the top *N* most highly-variable genes (HVGs) in the processed data for model training. This can be achieved with the *highly*_*variable*_*genes* function in *Scanpy* using the *cell*_*ranger* flavor.(46) This method was also used in the *scIB* benchmark to identify HVGs. We used *N* = 2.000.

A summary of the preprocessing pipeline is shown in Figure 2a. For the Baron, Segerstolpe, and Zheng68k datasets, only steps 5, 6, and 8 were performed, as these datasets are widely used for benchmarking cell type annotation, and their sample quality is assumed to be high. For the MacParland dataset, only steps 5 and 8 were performed as it comes pre-normalized. All four of these datasets are already annotated and step 7 was not required.

**Fig. 2.**
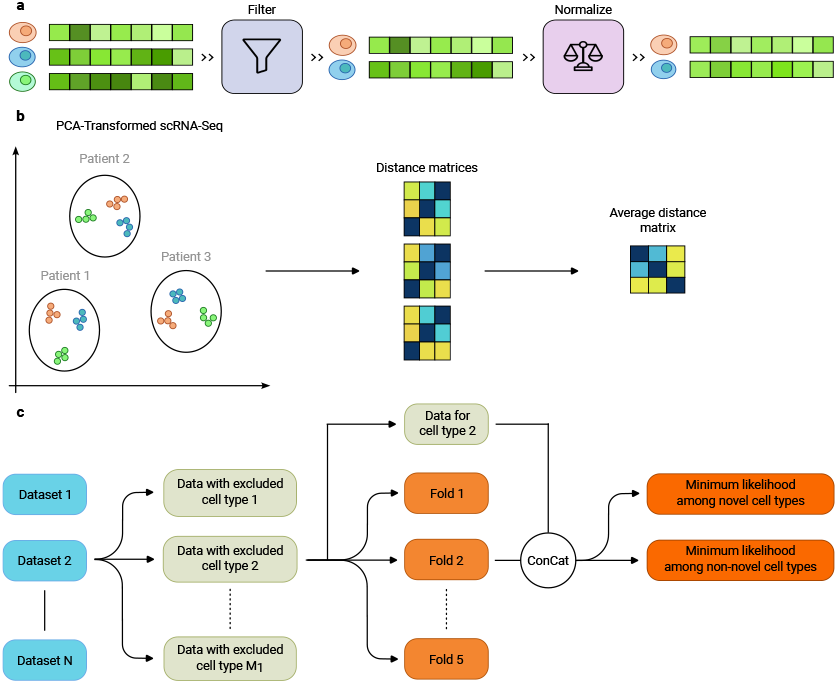
(a) Summary of the preprocessing pipeline, starting with scRNA-Seq data from multiple samples. The data are then filtered to ensure sample quality, and normalized to make samples comparable. (b) Overview of steps for calculating the cell type centroid loss (Equation 8). PCA is used to reduce the dimensionality of the scRNA-Seq data. The Euclidean distance matrix is calculated between cell type clusters in different batches; from this, the average distance matrix is calculated. (c) Framework for evaluating the likelihood thresholds for novel cell type detection. For each dataset, we begin by excluding one cell type at a time. Then, for each such data subset we use a 5-fold split for training/testing. The data of the excluded cell type is concatenated to the test fold, acting as a “novel” cell type. Finally, the minimum likelihood produced by CELLULAR is then aggregated for the non-novel and novel cell types separately.

### 3.4 Training

#### 3.4.1 Contrastive loss

Models were trained using CL. This is a technique where the model tries to produce outputs that are close to each other in space for augmented inputs of the same original input, called positive (+) input pairs, while being pushed away from all other inputs, called negative (−) input pairs.(47)

To apply CL, one needs to define a suitable loss function. The *normalized temperature-scaled cross entropy loss* (NT-Xent; Equation 1) is a loss function that is appropriate for CL when there are multiple negative inputs.(48, 49)

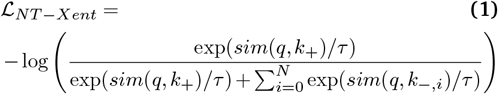

where *sim*() is the cosine similarity, *τ* represents the temperature parameter, *N* is the number of negative samples, *q* is the original input, *k* are augmented inputs where all *k*_*−,i*_ are negative inputs such that *q ≠ k*_*i*_, and *k*_+_ indicates the augmented version of the original input.

This learning strategy can be modified by instead stating that inputs belonging to the same label should strive to be as close to each other as possible while being far away from inputs of other labels, resulting in a semi-supervised approach. This is accomplished by modifying Equation 1 to be

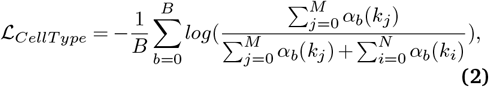

where *α*_*b*_ is defined as *α*_*b*_(*x*) = *exp*(*sim*(*q*_*b*_, *x*)*/τ*) and *q*_*b*_ is the current reference sample in the dataset, specified by the batch number *b* which consists of *B* samples. *k*_*j*_ are inputs of the same cell type as sample *q*_*b*_ while *q*_*b*_ ≠ *k*_*j*_ and *q*_*b*_ *≠k*_*i*_. *k*_*i*_ are inputs of all other cell type entries. *M* is the number of samples with the same label as *q*_*b*_ and *N* is the number of remaining samples, such that *M* + *N* = *B*. The loss calculation is performed for each sample *q*_*b*_ (i.e., for all data points in the batch), such that *ℒ*_*CellType*_ is an expectation value. This loss function is called the *soft nearest neighbor loss*.(50)

Using the loss described in Equation 2, one can enforce the creation of cell type clusters in latent space, but this loss alone does not necessarily reduce batch effects. To do so, we define an additional loss function

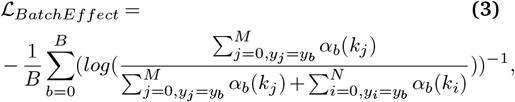

where *k*_*j*_ are inputs of the same batch effect key (e.g., laboratory) as sample *q*_*b*_ while *q*_*b*_ *≠k*_*j*_ and *q*_*b*_≠ *k*_*i*_. *k*_*i*_ are inputs of all other batches. *y*_*b*_ is the cell type of *q*_*b*_, *y*_*j*_ is the cell type of *k*_*j*_, and *y*_*i*_ is the cell type of *k*_*i*_, where *y*_*j*_ = *y*_*b*_ and *y*_*i*_ = *y*_*b*_ must be fulfilled. As in Equation 2, the total number of samples can be written as *B* = *M* + *N*. Using this loss we promote the dispersion of data points of the same batch within each cell type cluster.

ScRNA-Seq data tends to be unbalanced in terms of cell types and batch effects. Hence, it can be advantageous to weight the contributions of each label type to the loss differently. Weights can be calculated as

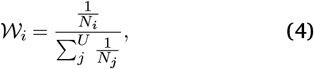

where *𝒲*_*i*_ is the weight to be applied to the loss of cell type label *i, N*_*i*_ is the number of samples of label *i* and *U* is the total number of unique cell types present in the data. By calculating the weights in this way, we promote less common cell types to have a larger impact during training, which counters the label unbalance. Weights are also calculated and used for batch effects in the same manner by redefining *N*_*i*_ to be the number of samples of batch effect *i* and *U* as the number of unique batch effect keys.

Combining Equations 2 and 4, we arrive at the final cell type loss

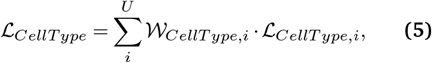

where *i* is a specific cell type and *U* is the number of unique cell types in the dataset. Similarly, for batch effects we get

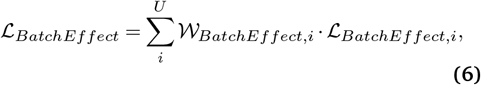

where *i* is a specific type of batch effect and *U* is the number of unique batch effect keys.

Finally, the two loss terms (Equations 5 and 6) are combined to arrive at the final contrastive loss

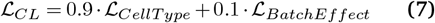

Here, the loss components are added together and weighted with 90% importance on the cell type loss and 10% on the batch effect loss, determined through empirical testing. A small batch effect loss is necessary to prevent excessive noise promotion but cannot be ignored completely; these percentages can be adjusted based on the project objectives.

#### 3.4.2 Adding a cell type centroid loss component

Enforcing a meaningful relativistic positioning of cell type clusters in the latent space can help the model learn biologically meaningful embeddings. To achieve this desired clustering behavior, an additional term, referred to in this study as the *centroid loss*, was added to the final loss function.

To derive the centroid loss, the first step is to transform scRNA-Seq expression levels using PCA into a latent space of the same dimension as the intended output of the model; in this study, this dimension is set to 100. PCA is used for its ability to extract dimensions responsible for significant variation in data, providing a lower-dimensional representation with reduced noise. From this latent space, we extract a Euclidean distance matrix containing distances between pairs of cell type cluster centroids (Figure 2b).

Assuming the main source of unwanted variation is due to the specified batch effect (patient ID in Figure 2b), we assume the distances between cell type clusters within each individual batch of data to be similar (see Section 4.1 for details). The average distance matrix across patients approximates the relative positioning of cell types in latent space. The relativistic distance matrix is obtained by normalizing the average centroid distance matrix by its maximum value. This is computed in both the PCA-transformed space and the model-produced space. Ideally, cell type clusters in the model’s latent space should preserve this relative positioning, enforced through the centroid loss term, defined as

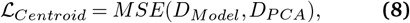

where *MSE* stands for mean squared error, *D*_*Model*_ is the relativistic centroid distance matrix produced by the model, and *D*_*P CA*_ is the same matrix but generated through dimensionality reduction of the scRNA-Seq via PCA. The workflow for calculating the centroid loss is further described in SI Section 3. Since not all cell types exist in each patient’s data, some pairwise distances may not be possible to calculate; these distances are masked when evaluating Equation 8.

Combining the CL loss (Equation 7) and the cell type centroid loss (Equation 8), we arrive at the final loss used in CELLULAR

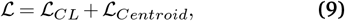

used to train all models developed in this work.

### 3.5 Evaluation

#### 3.5.1 Data integration benchmark

To evaluate the scRNA-Seq integration performance of CELLULAR and all its variations explored in this study, it was compared to other leading methods for omics integration using the *scIB* (14) suite of established benchmarks.

Performance of the models is evaluated in terms of (1) batch correction and (2) preservation of biological variation. This is evaluated using metrics such as *Normalized Mutual Information* (NMI), *Adjusted Rand Index* (ARI), *Average Silhouette Width* (ASW), *Isolated label Silhouette, Principal Component Regression* (PCR), *Graph Connectivity* (GC), *Isolated label F*_1_ *score*, and *Cell-cycle Conservation* (CC). All metrics are designed to fall within the range of 0 to 1, with 1 indicating better performance. We used the *scIB* package (14) for implementations of all metrics used in this study.

#### 3.5.2 Producing cell type representation vectors

A key application of the model’s embedding space is creating cell type representations. Since the model produces different embeddings for varying gene expression profiles of the same cell type, an aggregation method is needed to derive a single representation per cell type label. Possible approaches include: computing the centroid of each cluster, taking the median of each output dimension, or determining the medoid of each cluster. CELLULAR implements all three methods, allowing users to efficiently generate coherent cell type representations from scRNA-Seq data.

#### 3.5.3 Cell annotation

Another application of the learned embedding space is cell type classification. To explore this, a feed-forward neural network was trained using CELLULAR embeddings as input to predict cell types. The final layer applies a softmax function to generate likelihoods for each label, with training optimized via cross-entropy loss.(51) Since CELLULAR embeddings are generalizable, only the classifier weights are updated, while the encoder remains frozen. The model architecture is shown in Figure 1e.

Benchmarking of CELLULAR’s cell type annotation capabilities was done by comparing to the following models: scNym (18), Seurat (19), TOSICA (11), SciBet (20), and CellID (21). Three metrics were used to assess the performance of all models—accuracy, balanced accuracy, and F1 score—using *scikit-learn*.(52)

The models in this study include a built-in flag for automatic classifier hyperparameter optimization using *Optuna* (53). The data are split 80:20 for training and validation, and 100 *Optuna* trials are run to find optimal hyperparameters. Optimized hyperparameters include the number of neurons in the first and second hidden layers, initial learning rate, and dropout percentage.

#### 3.5.4 Novel cell type prediction

A cell type annotator can only predict cell types present in its training data. When applied to data containing novel cell types, it misclassifies them as known types, leading to incorrect predictions. However, novel cell types can still be detected as follows:

1. Predict cell type likelihood scores for each sample.
2. Min-max normalize likelihoods across all cell types, setting the maximum to 1.0 and the minimum to the inverse of the number of labels.
3. Extract the minimum likelihood from all predictions.
4. Compare this value to a predefined threshold (0.XX); values below this threshold suggest a novel cell type.

The intuition behind this approach is that if the model is uncertain about assigning any known label to a sample, it may indicate an unseen cell type. To investigate whether CELLULAR can be used for this purpose, an experiment was performed where one cell type was left out of training at a time from each of the MacParland, Baron, and Zheng68k datasets, followed by a 5-fold split. This eventually creates 5. 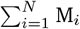 total folds, where M_*i*_ is the number of cell types in dataset *i* and *N* is the number of datasets. The data of the removed cell type is then concatenated to the test fold, so that one can make predictions both on known cell types and novel (excluded from training) cell types. For each test fold, dataset, and cell type left out during training, the minimum likelihood of any non-novel prediction was calculated along with the minimum likelihood of any novel guess (Figure 2c).

## 4 Results

### 4.1 Establishing patient ID as the main source of variability

During the formulation of the loss function described in Equation 8, we assume that patient ID is the main source of unwanted variation when calculating the Euclidean distances between cell type clusters. To investigate whether or not this is a reasonable assumption, we evaluated the coefficient of variation (*CV*) for each dataset:

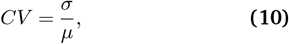

where *µ* is the mean Euclidean distance across patients between two cell type clusters and *σ* is the standard deviation of the Euclidean distance.

The CV score is a metric that can vary between 0 and infinity. A CV score of 1, for instance, means that the standard deviation equals the mean, indicating fairly large variability in the data. Ideally, one would observe CV scores equal to 0 for all pairs of cell type clusters, indicating that the normalized Euclidean distance between cell types *within each patient* is the same. However, in reality there exist many sources of error that can affect the CV calculation, and it is therefore unlikely to ever be 0, even if the underlying assumption is true. In the datasets explored in this work, the mean CV score tends to be around, and slightly below, 0.1, indicating a fairly low CV score for all datasets (Figure 3a). This supports the assumption that the major source of unwanted variability when calculating the Euclidean distances between cell type clusters in this work is indeed patient ID.

**Fig. 3.**
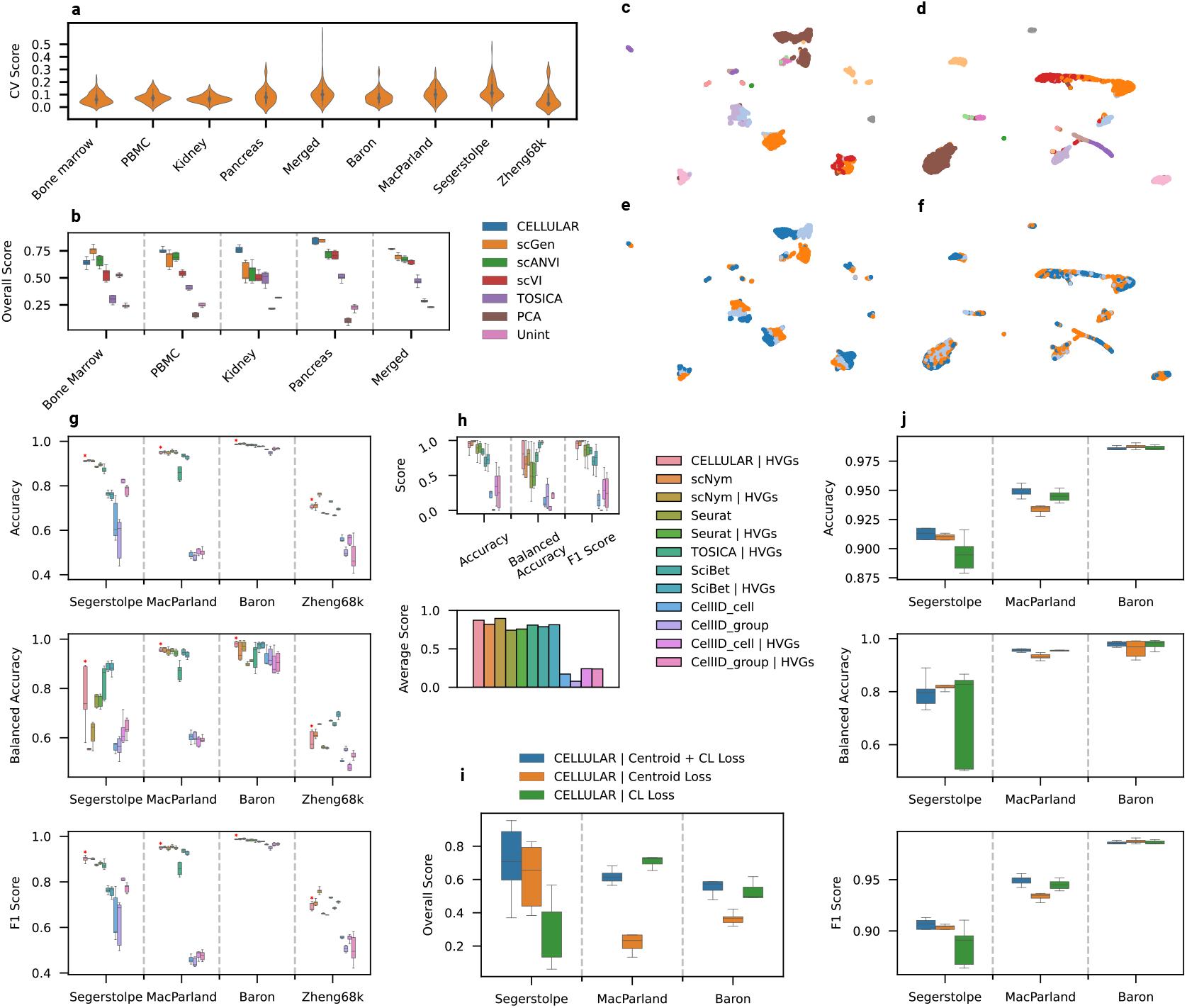
(a) Density of CV scores (Equation 10) for each dataset explored in this work. (b) Five-fold cross-testing on the bone marrow, PBMC, kidney, pancreas, and merged dataset, where the overall scores were calculated from min-max normalized *scIB*(14) metrics. The following models were benchmarked: CELLULAR, scGen, scANVI, scVI, TOSICA, PCA, and Unint (Unintegrated). The input to all methods were the top 2k HVGs. (c-f) UMAP of the first test fold for the kidney dataset when transformed by: (c) PCA, and colored by cell type, (d) CELLULAR, and colored by cell type, (e) PCA, and colored by patient ID, and (f) CELLULAR, and colored by patient ID. (g) Five-fold cross-testing on the Baron, MacParland, Segerstolpe, and Zheng68k datasets, where accuracy, balanced accuracy and F1 score are used as metrics. The following models were benchmarked: CELLULAR (emphasized by a red asterisk), scNym, Seurat, TOSICA, SciBet, and CellID. The input to the methods were either all genes or the top 2k HVGs. Legend in panel (h). (h) Box plots of min-max normalized mean metric values from panel (g) for the Baron, MacParland, Segerstolpe, and Zheng68k datasets. Below the box plots there is a bar plot of the average mean value of the metric scores for each model. (i) Five-fold cross-testing of CELLULAR on the Baron, MacParland, and Segerstolpe datasets, where the overall score was calculated from min-max normalized *scIB*(14) metrics. The three models benchmarked were CELLULAR using the centroid + contrastive loss, centroid loss, and the contrastive loss. The input to the models were the top 2k HVGs. (j) Five-fold cross-testing of CELLULAR using different losses on the Baron, MacParland, and Segerstolpe, datasets. The input to the models were the top 2k HVGs. Here, accuracy, balanced accuracy, and F1 score are shown. Legend in panel (i). In all box plots, the center line is the median and the box hinges the interquartile range, while the whiskers extend up to 1.5*×* the interquartile range.

### 4.2 Learning a generalizable scRNA-Seq embedding space

An ideal data integration algorithm would be able to enforce the biological variation in the data while minimizing variability caused by additional factors, such as patient ID, tissue, sequencing technology, etc. To investigate how well models performs on this task, we used the *scIB* package to benchmark models and evaluate their performance.(14) *scIB* computes various metrics, each of which we min-max normalize across methods for each metric and fold.

Here, we compared multiple existing methods to CELLULAR. As a control, we also fed the input data directly to the benchmark to obtain the corresponding metrics; we refer to this approach as “Unint” (short for “Unintegrated”) throughout the study. We benchmarked all models on the bone marrow, PBMC, kidney, pancreas, and merged datasets. In Figure 3b we see how *scIB* ranks CELLULAR among the best performing models for each dataset. The performance metrics for all models and for each dataset can be explored in more detail in SI Sections 2.1–2.5.

To get a visual understanding of how CELLULAR embeds different cell types, UMAP visualizations were prepared from one of the test folds of the kidney dataset following transformation with PCA (Figures 3c,e) and CELLULAR (Figures 3d,f), respectively. What we observe is a significant reduction in batch effects when using CELLULAR for embedding the data versus when using PCA; the data points are better integrated within and across clusters in Figures 3d&f relative to the clusters in Figures 3c&e.

### 4.3 Cell type annotation

To demonstrate a downstream use case for the cellular embeddings learned by CELLULAR, we used the learned embeddings for the most ubiquitous classification task in molecular biology: cell type annotation. Using four popular datasets (Segerstolpe, MacParland, Baron, and Zheng68k), we compared the accuracy, balanced accuracy, and F1 score of our annotator to multiple other established models (Figure 3g).

We found that CELLULAR was among the top performing methods for cell type annotation across all datasets and metrics explored. No single method, however, consistently performed best overall. To estimate the “overall performance,” we computed for each model the mean metric for each dataset and min-max normalized the means across models for each dataset (to make the metrics comparable between datasets); the results are visualized in Figure 3h using a box plot.

Balanced accuracy is useful for evaluating scRNA-Seq annotation models as it accounts for class imbalance, giving more weight to common cell types, whereas ordinary accuracy treats all labels equally. While balanced accuracy may be a more informative metric, we averaged all metrics equally, computing the arithmetic mean of accuracy, balanced accuracy, and F1 score. The resulting “Average Score” (Figure 3h) provides a clearer performance comparison, showing scNym as the top model with highly variable genes (HVGs), followed closely by CELLULAR. Nevertheless, we note that in the original scNym study (18) the authors did not explore the use of HVGs; this is our contribution, and without HVGs, scNym would not be among the top performers for cell type annotation. In general, we observed that the use of HVGs, rather than all genes, led to better performance for all models benchmarked.

### 4.4 Loss function analysis

CELLULAR uses a loss function specifically engineered for creating as generalizable an embedding space as possible from scRNA-Seq data (Equation 9). As an ablation study, we evaluate the performance of CELLULAR with each of the two loss components individually, namely the contrastive (Equation 7) and centroid (Equation 8) loss terms. This comparative analysis (Figure 3j) underscores the importance of using the complete loss function.

We observed that using the cell type centroid loss alone (Equation 8) led to weak performance on the MacParland dataset, while using the contrastive loss alone (Equation 7) led to weak performance on the Segerstolpe dataset. The best performance on the accuracy, balanced accuracy, and F1 score (Figure 3j) was always observed when using the complete loss function for all three datasets explored (Segerstolpe, MacParland, and Baron). Similarly, the best performance on the overall score from the *scIB* benchmark was observed when using the complete loss function (Figure 3i).

### 4.5 Novel cell type detection

As seen in the previous sections, CELLULAR demonstrates promising performance for creating a generalizable embedding space for cell types, as well as good performance in cell type classification. An even more challenging application of the model would be to use it for novel cell type detection, i.e., identifying cell types that don’t fit into any of the classes that the model has previously seen.

When it comes to classification tasks, it is well-known that neural networks generalize poorly to classes not seen during training due to their reliance on learned patterns from the training data, thus limiting their ability to recognize novel patterns corresponding to new classes. This limitation can be leveraged for novel cell type detection by setting a likelihood threshold for known classes. If a sample falls below this threshold across all classes, it is considered novel. In other words, if the model lacks confidence in assigning a known label to a given gene expression pattern, the pattern may correspond to a new cell type. Here, ‘new’ or ‘novel’ refers to being unseen by the model, not necessarily biologically novel, though confirming a truly new cell type would be the ultimate validation of this approach.

We investigated how picking different threshold values influenced the predictions made by the novel cell type detection model, and found that a likelihood threshold of 0.25 was generally acceptable (Figure 4a). We found that this value led to high precision (0.81) while maintaining reasonable coverage (0.32). The impact of the threshold value on precision and coverage is further illustrated in Figure 4b. Our findings suggest that one can vary the threshold value for novel cell type detection depending on whether prioritizes precision or coverage. Here, 0.25 reflects a reasonable compromise between coverage and precision, slightly favoring precision to avoid the prediction of false positives.

**Fig. 4.**
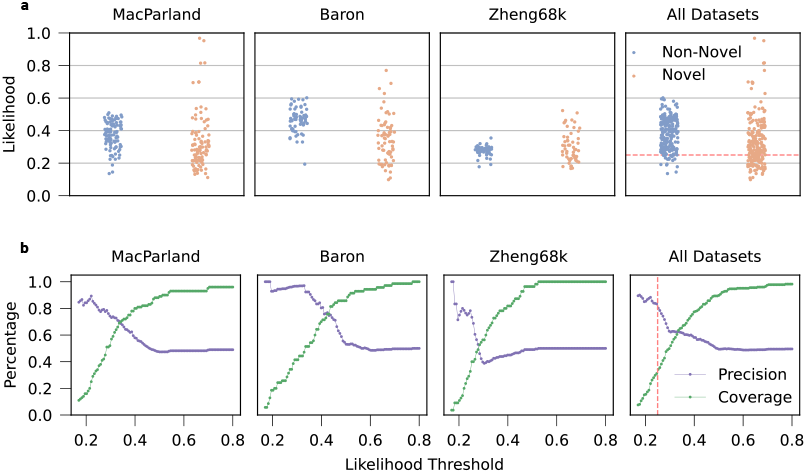
(a) Jitter plot visualizing the distribution of the minimum likelihood attained by predictions made using CELLULAR for both non-novel and novel cell types. Each point represents the minimum likelihood observed for predictions of observed and unobserved cell types when one cell type was omitted from training across various test folds and datasets (MacParland, Baron, Zheng68k, and All Datasets). The red dotted line represents a likelihood threshold of 0.25; all data points falling below the line would be novel cell types according to the model. The *All Datasets* panel combines data from MacParland, Baron, and Zheng68k into a single plot. To ensure comparability, data points are min-max normalized based on each dataset’s likelihood range. (b) Line plot visualizing the relationship between the likelihood threshold used and the precision and coverage for detecting novel cell types using CELLULAR. The red dotted line represents a likelihood threshold of 0.25, which leads to a precision of 0.81 and a coverage of 0.32 for novel cell type detection across all datasets. A trade-off between precision and coverage is evident, providing the option to choose thresholds based on desired model behavior.

## 5 Discussion

Our model displays impressive performance on two benchmarks. Of the diverse models benchmarked herein, CELLULAR was overall the best performing model on scRNA-Seq data integration, and second best for cell type classification. On top of this, CELLULAR also has the ability to identify the presence of novel cell types in data. Its robust performance makes it a valuable tool for both bioinformaticians and DL researchers in single-cell data analysis.

### 5.1 The importance of HVGs

The best performing model for cell classification in this work was *scNym*, but it only outperforms CELLULAR when the input data are trimmed down to the top 2k HVGs, something not investigated in the original study.(18) In fact, the use of HVGs improved the performance of all models benchmarked in this study. As such, we want to emphasize the importance of high-quality inputs to DL methods for scRNA-Seq data integration, as these methods are quite sensitive to how the raw data are processed.

Researchers working on the development of new methods for cell annotation and related tasks with scRNA-Seq data should consider limiting the model input data to only HVGs to promote better generalization and utility in their tools.

### 5.2 Novel cell type detection

Detecting novel cell types is inherently challenging. In the Zheng68k dataset (Figure 4b), high precision comes at the cost of coverage, making the model prone to missing novel cell types. In contrast, the Baron dataset achieves both high precision and coverage, highlighting the model’s dependence on dataset characteristics. This variability complicates optimal threshold selection. A 0.25 threshold, set by aggregating data across datasets, yielded high precision with acceptable coverage, though its generalizability beyond these datasets remains untested. Thus, the model is best used as a supplementary verification tool in novel cell type detection pipelines. CELLULAR can boost confidence when detecting novelty but cannot guarantee the absence of novel cell types when it fails to do so.

### 5.3 Possible data augmentation strategies

CELLULAR’s performance may be further improved through data augmentation strategies. Possible augmentations for scRNA-Seq data include randomly masking a specified percentage of gene expressions (i.e., making them zero) and adding Gaussian noise. The former aims to simulate the occurrence of errors during sequencing, leading to the dropout of gene expression. The latter can be done by randomly replacing a specified percentage of gene expressions with a value drawn from a Gaussian distribution fit to the expression distribution of each gene in a given cell type.

Since gene expression is often dependent on other genes, randomly removing expression levels may hinder the model’s ability to learn these connections, making this augmentation method potentially undesirable, though it could also improve generalization. Further, adding Gaussian noise to gene expression levels could be used to generate artificial training data. However, this assumes minimal noise within each cell type, which is a potentially strong assumption. Investigating both strategies in future work could provide valuable insights.

### 5.4 Future applications

#### 5.4.1 Cell type representations

The core strength of CELLULAR is its ability to generate a generalizable embedding space, enabling various downstream tasks like cell type annotation and novel cell type detection. Another implemented but unexplored application is the generation of cell representation vectors for each unique cell type.

This was achieved using three methods: calculating the centroid, median, or medoid of each cell type cluster. However, their effectiveness in downstream applications remains untested and is left for future research. Potential uses include identifying quantitative structure-activity relationships (QSAR) conditioned on cell type or integrating cell type information into molecular generative models for drug discovery—capabilities largely absent in data-driven drug discovery literature.

Additionally, future work could explore CELLULAR’s ability to extract broader biological insights by applying it to classification and clustering tasks for drug or gene perturbations after training solely on cell types.

#### 5.4.2 Multi-modal DL approaches

In the future, this approach can be integrated with AI models trained on cell assay data to develop multimodal cell representation learning models.

CL can also be leveraged to learn a joint embedding space from scRNA-Seq data and cell images, such as Cell Painting images. Cell Painting is a highcontent, image-based assay that captures cellular state information.(54) For example, it has been shown that Cell Painting images can distinguish cell health under various CRISPR perturbations designed to target genes linked to biological pathways, often inducing morphological changes.(55) These images serve as an unbiased, quantitative representation of cell conditions.(56)

Integrating Cell Painting with scRNA-Seq data could enhance CELLULAR’s latent space by incorporating additional biological context, improving its generalizability and downstream utility. However, this approach requires a large number of Cell Painting images corresponding to cell types with available scRNA-Seq data, which remains a major limitation. Due to the absence of a public Cell Painting database covering diverse cell types relevant to this study, this direction was not pursued further. Nevertheless, it remains an intriguing avenue for future research.

Yet another way of modifying CELLULAR would be to introduce other forms of omics data, such as single-cell assay for transposase-accessible chromatin sequencing (scATAC-Seq) data. With CL, one version of CELLULAR could learn from scRNA-Seq while another learns from scATAC-Seq. Integrating multiple data types—scRNASeq, scATAC-Seq, and Cell Painting images—could enrich the latent space by leveraging complementary biological information and improving generalizability. This multi-modal approach may provide a more accurate representation of cell type relationships and distinctions.

Just as multi-modal deep learning revolutionized text-conditioned image generation, its application in molecular biology could deepen our understanding of cellular development.

### 5.5 Limitations

As with most models, CELLULAR has a few limitations. One limitation is the time required for training, which presents a problem if one wants to train a model using a large number of cells. The biggest dataset used for training in this study was the Zheng68k dataset, containing around 68k cells. While it was manageable to train with this amount of data (∼ 30 minutes to train the encoder, ∼100 minutes to train the classifier, for a dataset of ∼50k cells), larger datasets containing on the order of millions of cells could present a challenge. The relatively slow training is due to the complex loss function used by CELLULAR. A large amount of computations are needed per data point. Developing methods to reduce the computational complexity of the model could thus enhance its scalability and applicability to larger datasets.

Secondly, the performance of the benchmarked models, including CELLULAR, is highly dataset-dependent, with model rankings varying significantly across datasets. To mitigate this, we used a diverse set of datasets spanning different tissues and laboratories. However, these datasets represent only a fraction of what is possible with scRNA-Seq and other omics technologies. Researchers should exercise caution when generalizing conclusions to datasets that differ significantly from those used in this study.

Finally, the cell type centroid loss requires prior annotation for training but is only used to structure the embedding space, not during inference. This could be a limitation for datasets where prior annotation is challenging or unreliable. While labels refine the embedding space during training, the model does not automatically assign cell type labels to new data.

## 6 Conclusion

In this study, we present a robust new method for leveraging scRNA-Seq data to learn a generalizable embedding space for cells using CL. The model, CELLULAR, is effective at mitigating batch effects while enforcing biologically relevant information. The approach was developed into an open-source Python package, with multiple downstream applications, including cell type annotation and novel cell type detection, also integrated into the package. This comprehensive approach to cell representation learning can empower other researchers to explore this method for learning meaningful cellular embedding spaces from other single-cell transcriptomics datasets.

This work advances the field of bioinformatics via the following key contributions:

- We introduce a new model for cell representation learning from scRNA-Seq data, and demonstrate that it outperforms the current leading methods in data integration. The model also demonstrates robust performance on cell type annotation and novel cell type detection tasks.
- By using a carefully crafted contrastive loss, we demonstrate that our model is robust to batch effects while preserving biologically relevant information.
- We show how pruning preprocessed scRNA-Seq data to only the most highly variable genes (HVGs) improves their utility for cell annotation, regardless of the model or dataset used.

Finally, we offer suggestions for future directions in cell representation learning using this approach, highlighting the significant promise we see in multi-modal DL, particularly with cell images and other sc-omics methods.

## Supplementary information

In the Supplementary Information (SI) we provide: UMAP visualizations for the various datasets used in this work; additional details on the embedding space benchmarks for all datasets; pseudo-code for the centroid loss algorithm; and attributions.

## Acknowledgements

The authors acknowledge computing resources from the National Academic Infrastructure for Supercomputing in Sweden (NAISS), partially funded by the Swedish Research Council through grant agreement no. 2022-06725. RM thanks the Wallenberg AI, Autonomous Systems, and Software Program (WASP) for funding.

## Author contributions

RM conceived and supervised the study. RM and LA discussed and designed the models developed in this study, and designed experiments together. LA carried out all data collection, data preprocessing, model training, and computational analysis. LA wrote all code presented in this study. Both authors discussed the results. LA prepared an initial draft of the manuscript, and both authors contributed to the writing and final structure of the manuscript.

## Data availability

CELLULAR has been implemented as a stand-alone Python package, available at: github.com/LeoAnd00/CELLULAR. Further, the reproducibility code for this work is available at: github.com/LeoAnd00/CELLULAR_reproducibility.

The public data used for this work is available at: doi.org/10.5281/zenodo.12169799.

