## Supplementary Information for "Contrastive Learning for Robust Cell Annotation and Representation from Single-Cell Transcriptomics"

February 27, 2025

**3 Centroid loss algorithm** **16**

**4 Attributions** **17**

### 1 Data preprocessing

#### 1.1 Bone marrow dataset

UMAP of the bone marrow dataset showing all 9 cell types (Figure S1a) and 4 patient IDs / sequencing methods (Figure S1b). The batch effects are not that severe for this dataset.

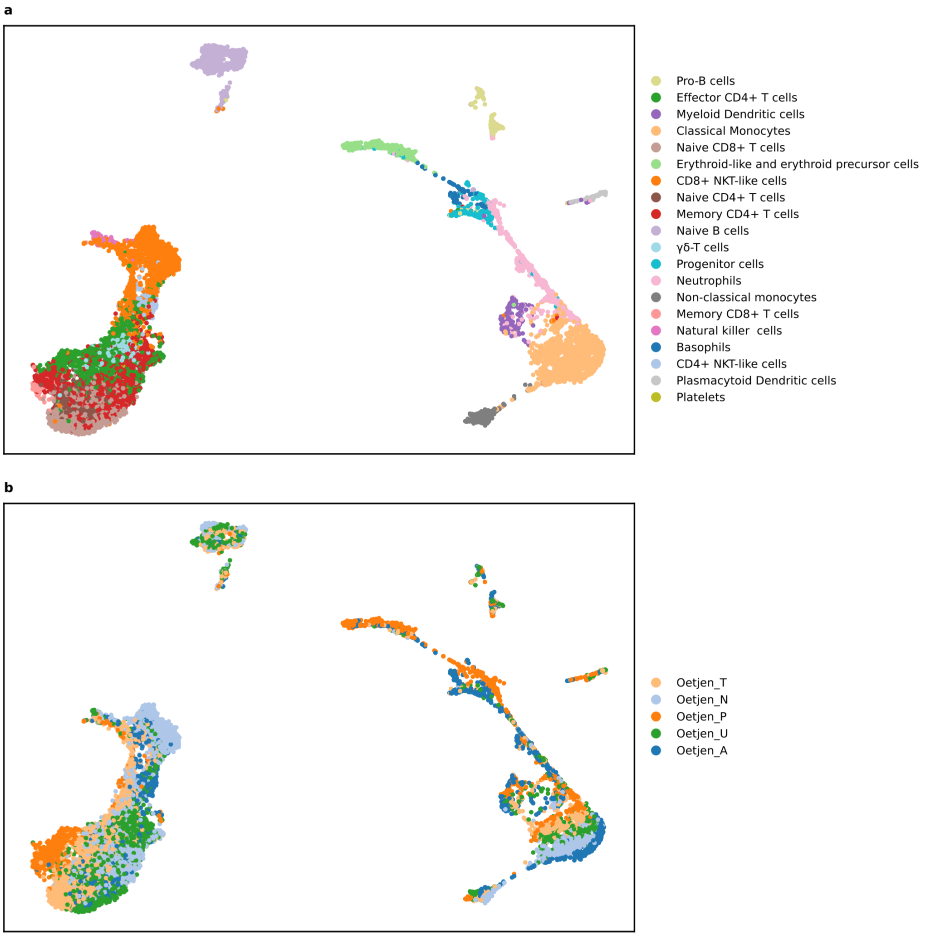

Figure S1: **a** UMAP visualization of cell type clusters in the bone marrow dataset. **b** UMAP highlighting batch effect caused by patient ID method.

#### 1.2 PBMC dataset

UMAP of the PBMC dataset showing all 9 cell types (Figure S2a) and 4 patient IDs / sequencing methods (Figure S2b). The batch effects are much more severe for this dataset compared to the bone marrow dataset.

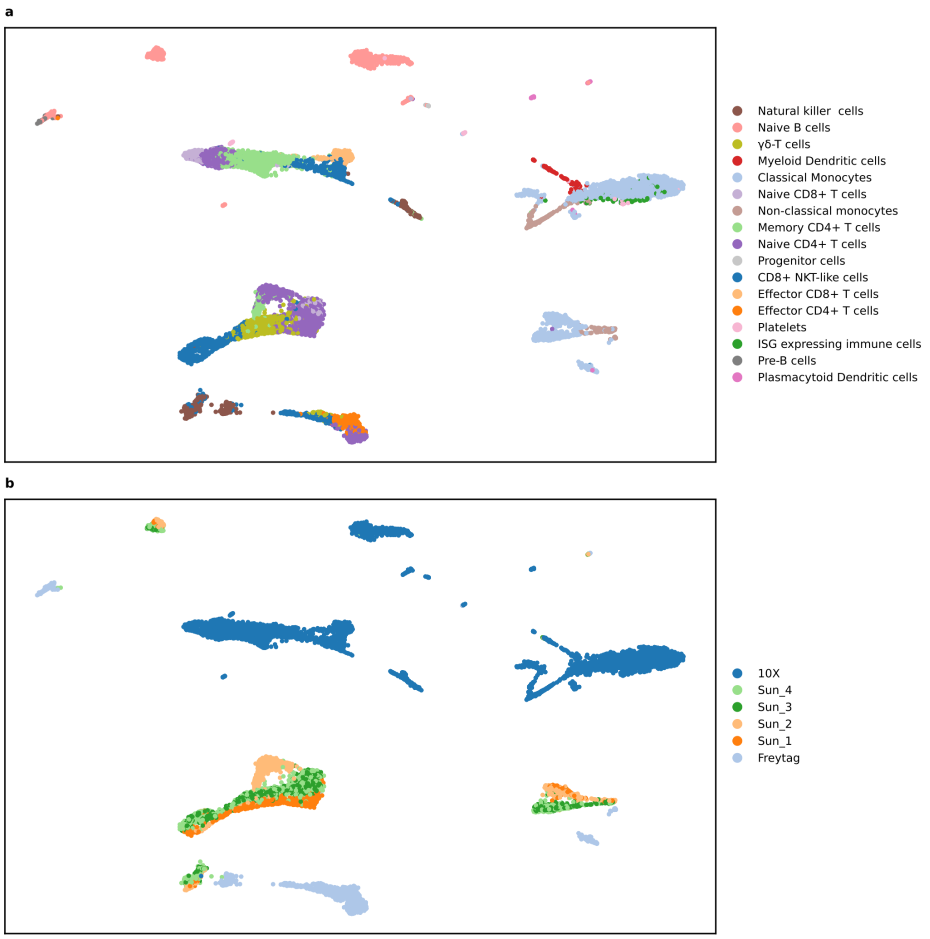

Figure S2: **a** UMAP visualization of cell type clusters in the PBMC dataset. **b** UMAP highlighting batch effect caused by patient ID method.

#### 1.3 Pancreas dataset

UMAP of the pancreas dataset showing all 9 cell types (Figure S3a) and 4 patient IDs / sequencing methods (Figure S3b). The batch effects are very severe for this dataset.

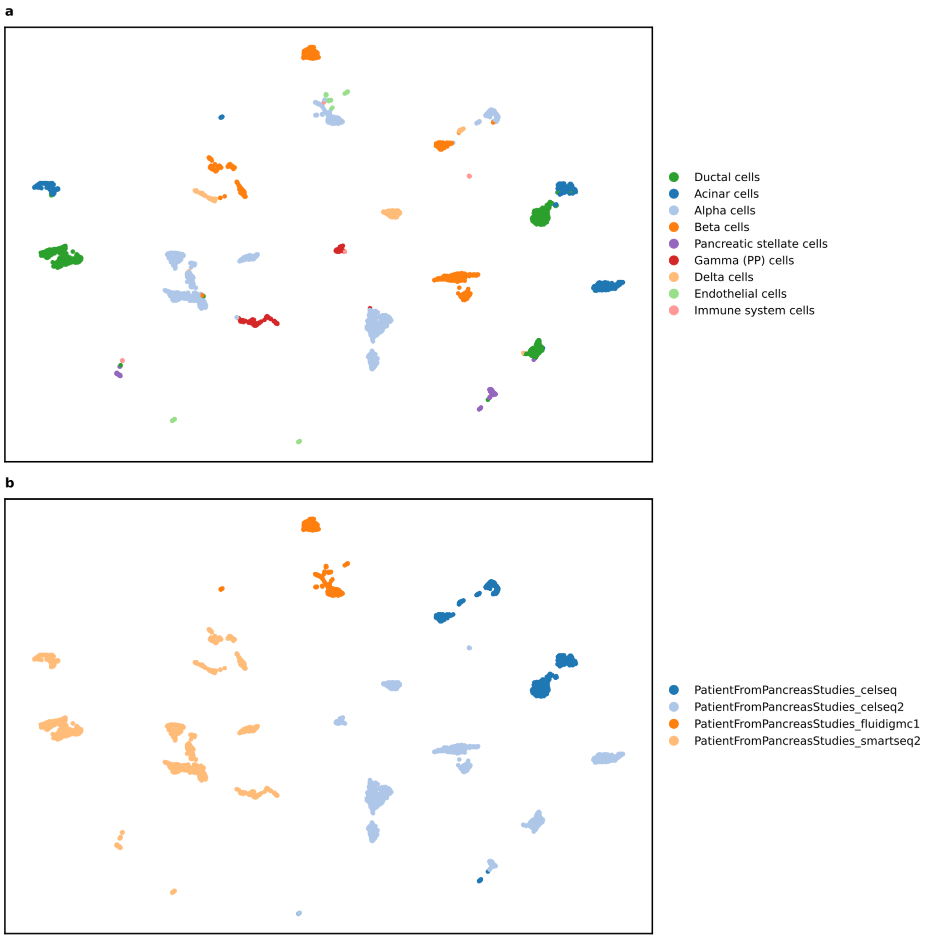

Figure S3: **a** UMAP visualization of cell type clusters in the pancreas dataset. **b** UMAP highlighting batch effect caused by patient ID / sequencing method.

#### 1.4 Kidney dataset

UMAP of the kidney dataset showing all 15 cell types (Figure S4a) and 3 patient IDs (Figure S4b). There are visible batch effects.

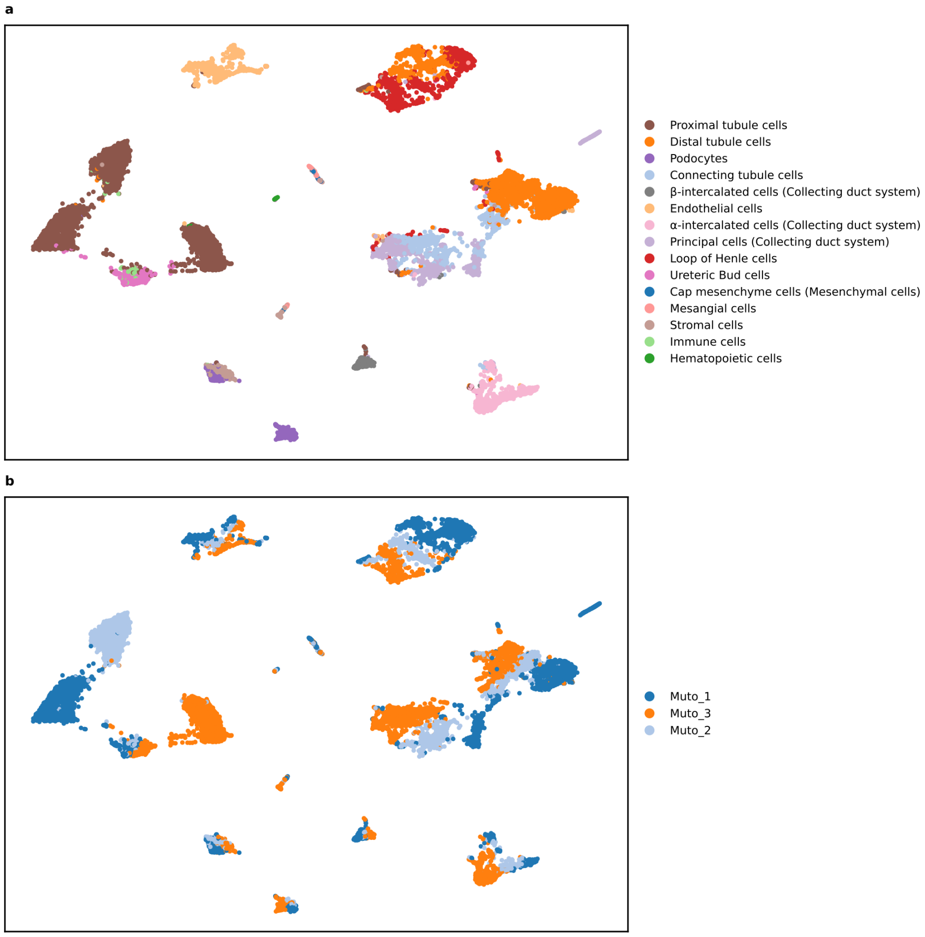

Figure S4: **a** UMAP visualization of cell type clusters in the kidney dataset. **b** UMAP highlighting batch effect cause by patient ID.

#### 1.5 Merged dataset

The entire merged dataset was visualized via UMAP and colored according to all 46 cell types (Figure S5a) and 18 batch effect elements (Figure S5b).

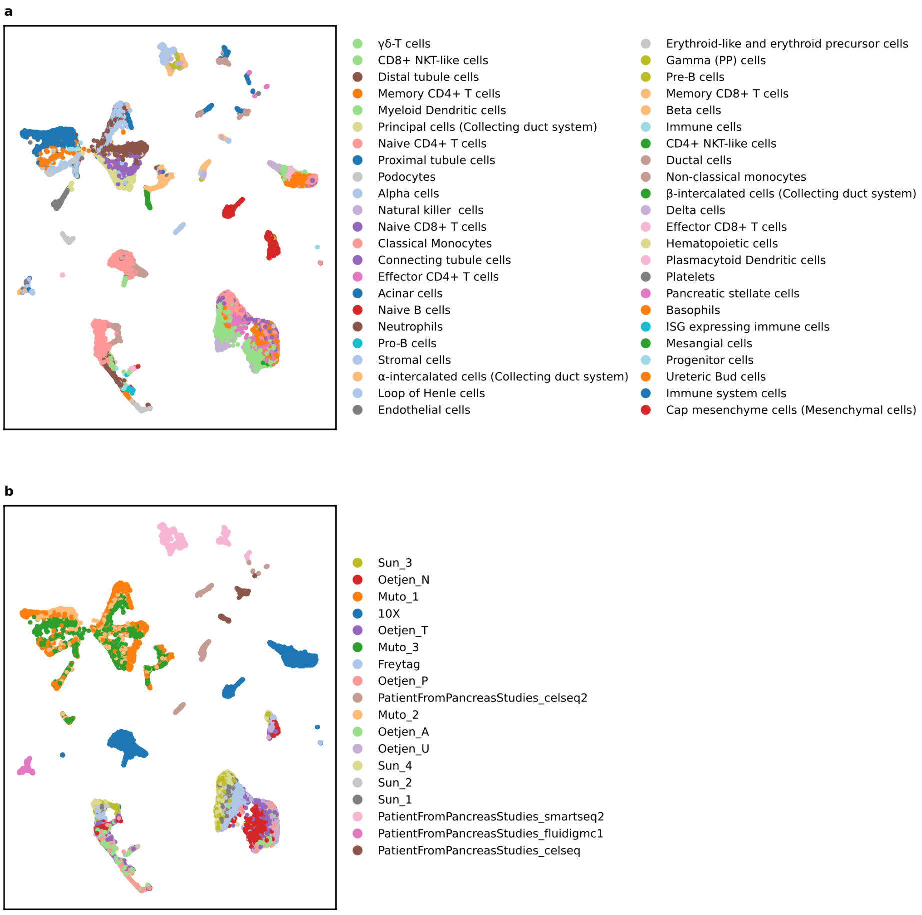

Figure S5: **a** UMAP visualization of cell type clusters in the full merged dataset. **b** UMAP highlighting the batch effects when merging the bone marrow, PBMC, kidney, and pancreas datasets.

#### 1.6 Baron dataset

UMAP of the Baron dataset showing all 14 cell types (Figure S6a) and 4 patient IDs (Figure S6b).

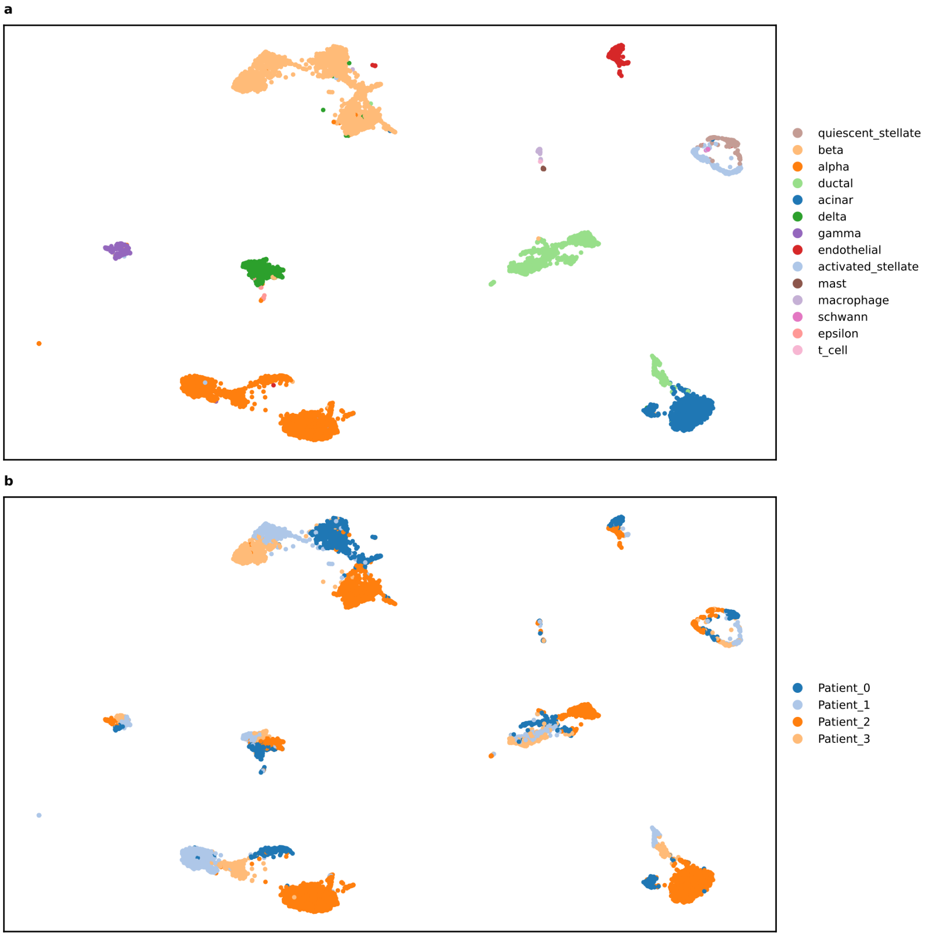

Figure S6: **a** UMAP visualization of cell type clusters in the Baron dataset. **b** UMAP highlighting batch effect cause by patient ID.

#### 1.7 MacParland dataset

UMAP of the MacParland dataset showing all 20 cell types (Figure S7a) and 5 patient IDs (Figure S7b).

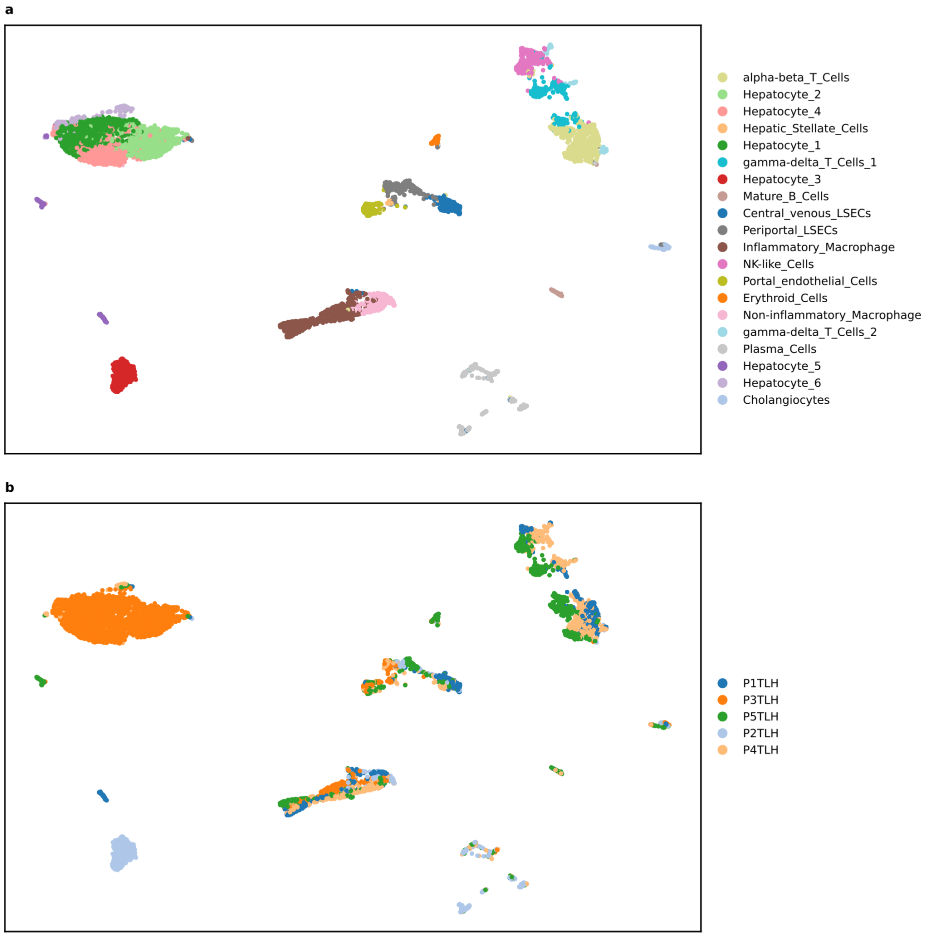

Figure S7: **a** UMAP visualization of cell type clusters in the MacParland dataset. **b** UMAP highlighting batch effect cause by patient ID.

#### 1.8 Segerstolpe dataset

UMAP of the Segerstolpe dataset showing all 15 cell types (Figure S8a) and 10 patient IDs (Figure S8b).

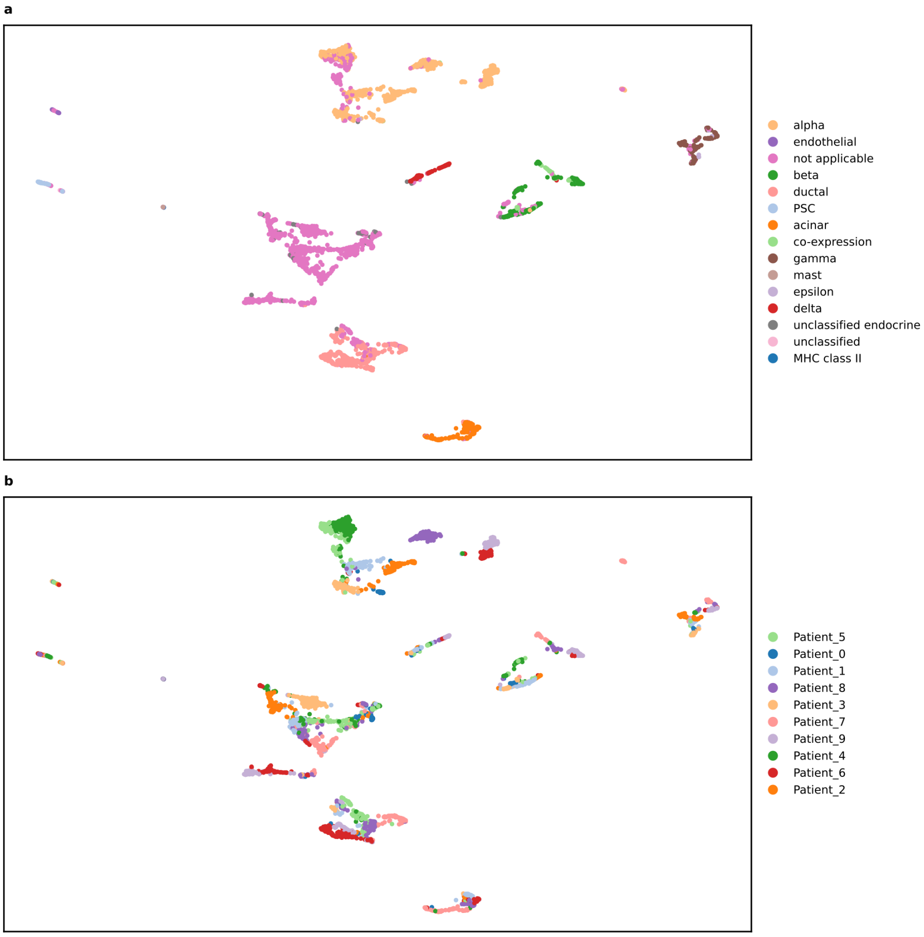

Figure S8: **a** UMAP visualization of cell type clusters in the Segerstolpe dataset. **b** UMAP highlighting batch effect cause by patient ID.

#### 1.9 Zheng68k dataset

UMAP of the Zheng68k dataset showing all 11 cell types (Figure S9a) and 8 unique barcode IDs (Figure S9b).

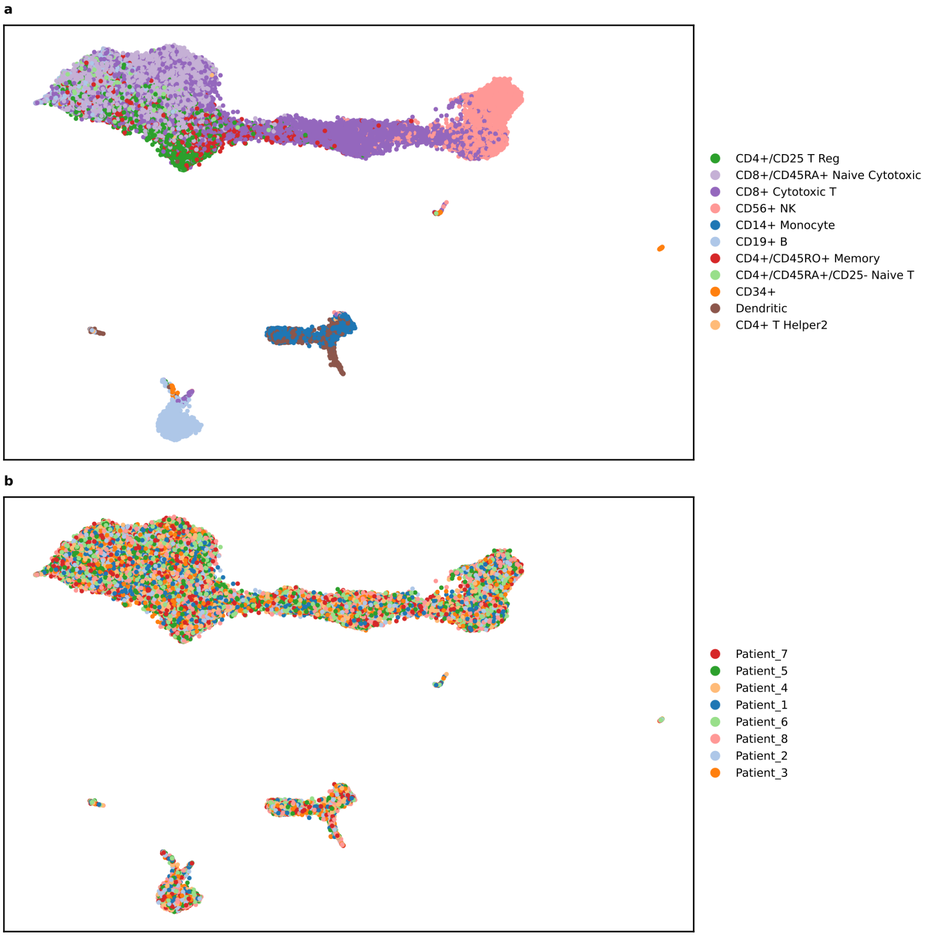

Figure S9: **a** UMAP visualization of cell type clusters in the Zheng68k dataset. **b** UMAP highlighting batch effect cause by the barcode IDs.

### 2 Generalizable scRNA-Seq embedding space benchmark

#### 2.1 Bone marrow dataset

The results for all 12 metrics, including overall bio, overall batch, and overall scores for all models when trained on the bone marrow dataset are shown below.

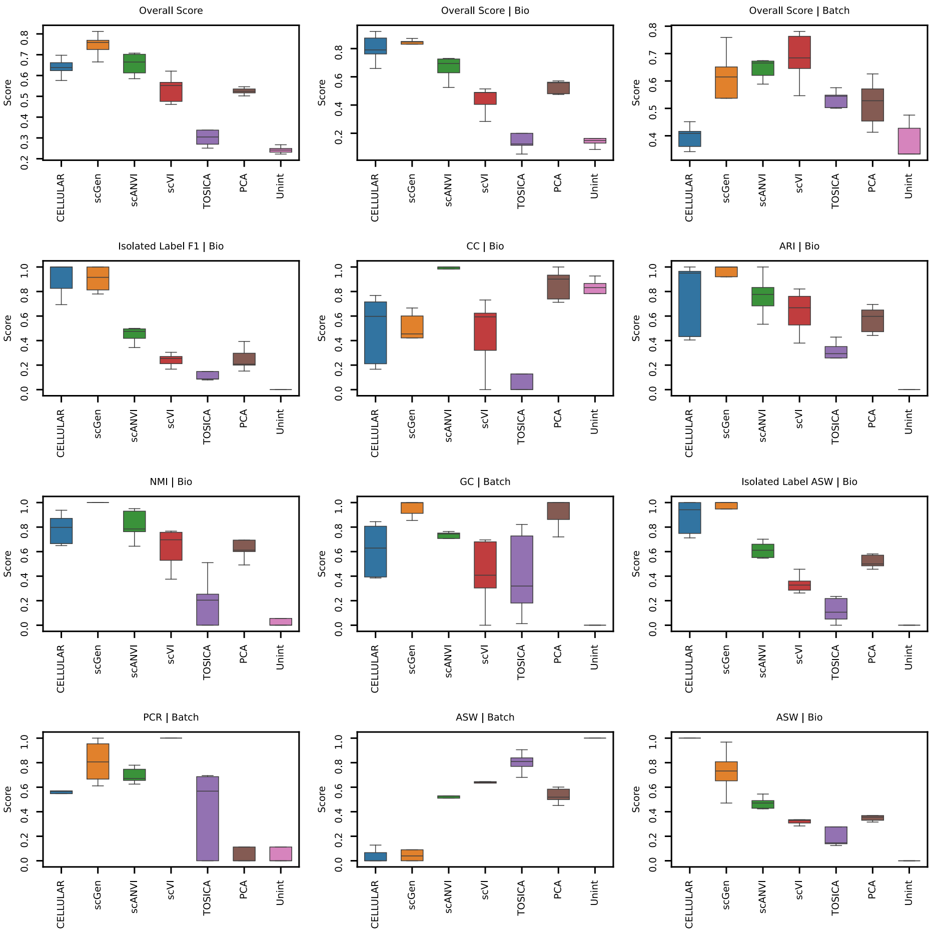

Figure S10: Five-fold cross-testing on the bone marrow dataset, where all metrics are calculated. The following models are used in the benchmark: Unint (Unintegrated), PCA, TOSICA[1], scVI[2], scANVI[3], scGen[4], and CELLULAR. The input to the methods are the top 2k HVGs. Box plots show the median displayed as the center line, the interquartile range as the box hinges, and the whiskers extending up to 1.5 times the interquartile range.

#### 2.2 PBMC dataset

The results for all 12 metrics, including overall bio, overall batch, and overall scores for all models when trained on the PBMC dataset are shown below.

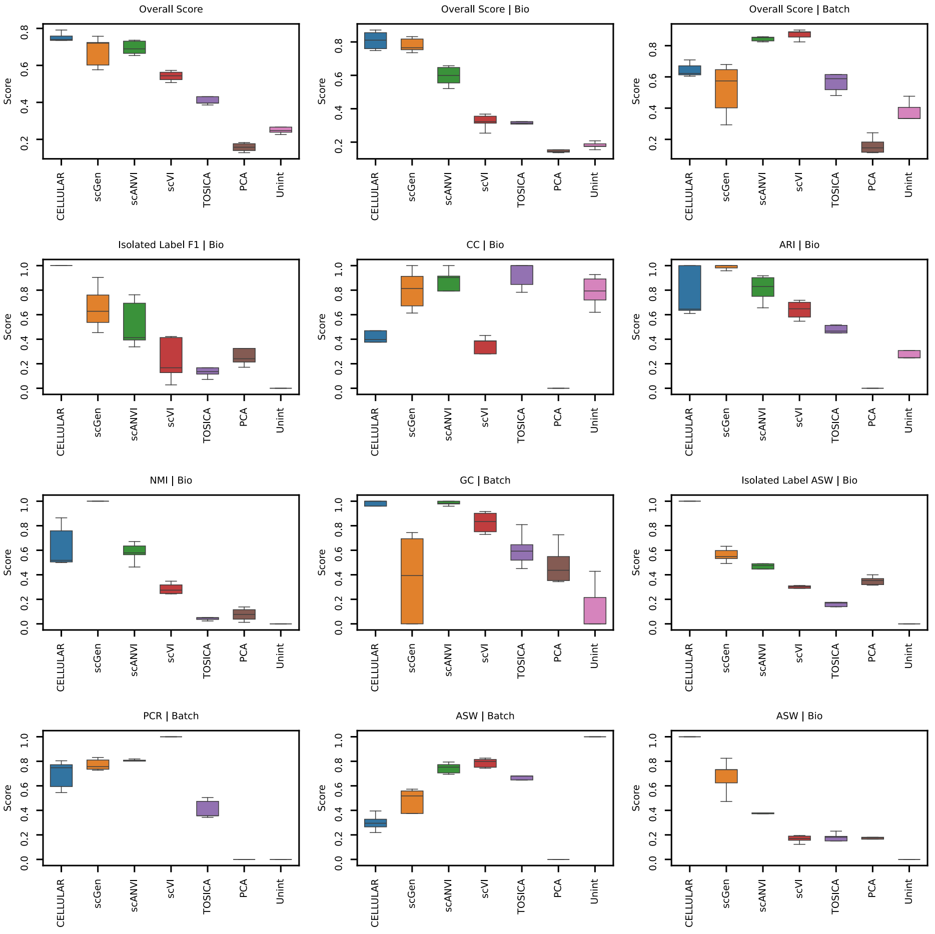

Figure S11: Five-fold cross-testing on the PBMC dataset, where all metrics are calculated. The following models are used in the benchmark: Unint (Unintegrated), PCA, TOSICA[1], scVI[2], scANVI[3], scGen[4], and CELLULAR. The input to the methods are the top 2k HVGs. Box plots shows the median displayed as the center line, the interquartile range as the box hinges, and the whiskers extending up to 1.5 times the interquartile range.

#### 2.3 Kidney dataset

The results for all 12 metrics, including overall bio, overall batch, and overall scores for all models when trained on the kidney dataset are shown below.

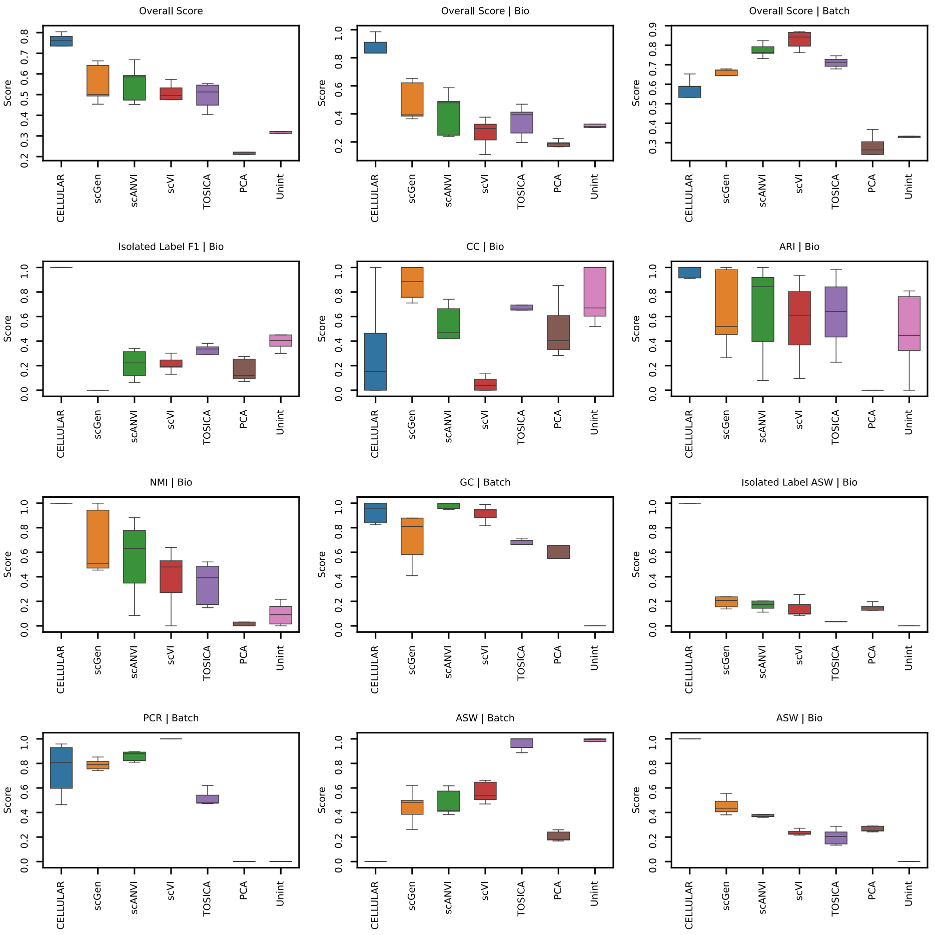

Figure S12: Five-fold cross-testing on the kidney dataset, where all metrics are calculated. The following models are used in the benchmark: Unint (Unintegrated), PCA, TOSICA[1], scVI[2], scANVI[3], scGen[4], and CELLULAR. The input to the methods are the top 2k HVGs. Box plots shows the median displayed as the center line, the interquartile range as the box hinges, and the whiskers extending up to 1.5 times the interquartile range.

#### 2.4 Pancreas dataset

The results for all 12 metrics, including overall bio, overall batch, and overall scores for all models when trained on the pancreas dataset are shown below.

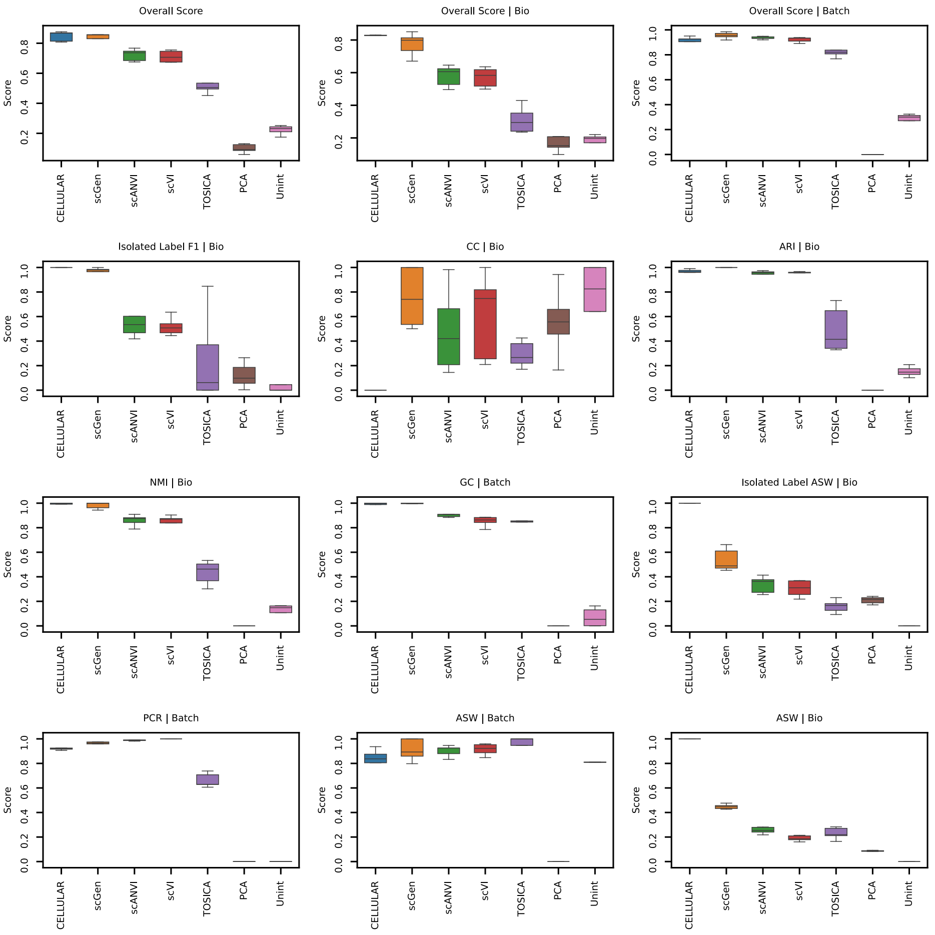

Figure S13: Five-fold cross-testing on the pancreas dataset, where all metrics are calculated. The following models are used in the benchmark: Unint (Unintegrated), PCA, TOSICA[1], scVI[2], scANVI[3], scGen[4], and CELLULAR. The input to the methods are the top 2k HVGs. Box plots shows the median displayed as the center line, the interquartile range as the box hinges, and the whiskers extending up to 1.5 times the interquartile range.

#### 2.5 Merged dataset

The results for all 12 metrics, including overall bio, overall batch, and overall scores for all models when trained on the merged dataset are shown below.

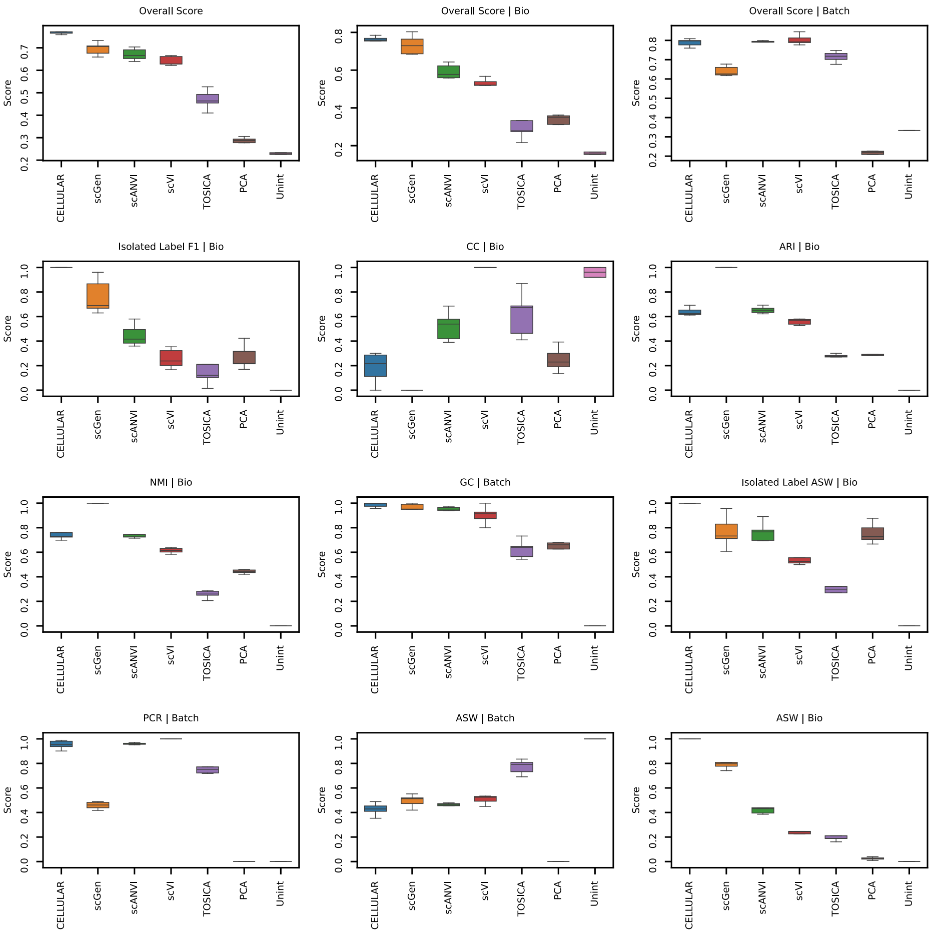

Figure S14: Five-fold cross-testing on the merged dataset, where all metrics are calculated. The following models are used in the benchmark: Unint (Unintegrated), PCA, TOSICA[1], scVI[2], scANVI[3], scGen[4], and CELLULAR. The input to the methods are the top 2k HVGs. Box plots shows the median displayed as the center line, the interquartile range as the box hinges, and the whiskers extending up to 1.5 times the interquartile range.

### 3 Centroid loss algorithm

Below, Algorithm 1 outlines of the algorithm for calculating the cell type centroid loss introduced in Section 3.4.2 (note that the algorithm is split over two pages).

**Algorithm 1** Cell Type Centroid Loss Algorithm

*// Step 1: Reference Distance Matrix*
Input: *adata // Annotated data object*
Define: $X = adata.X$ *// scRNA-Seq data*
Define: $X_{PCA} = PCA(X, dim = 100)$
Define: $A =$ "*cell_type*"; $B =$ "*patientID*"
Define: $unique_A = unique(adata.obs[A])$
Define: $unique_B = unique(adata.obs[B])$
Define: $centroids = \{\}$
**for** $\beta$ in $unique_B$ **do**
    **for** $\alpha$ in $unique_A$ **do**
        Define mask: $mask = (adata.obs[B] == \beta)$ & $(adata.obs[A] == \alpha)$
        Apply mask and calculate centroid:
$centroids[(\beta, \alpha)] = mean(X_{PCA}[mask], axis = 0)$
    **end for**
**end for**
Initialize distance matrix:
$D_{ref} = zeros(len(unique_A), len(unique_A))$
**for** $\alpha_1$ in $unique_A$ **do**
    **for** $\alpha_2$ in $unique_A$ **do**
        $\delta = []$
        **for** $\beta$ in $unique_B$ **do**
            $\delta_\beta = EuclideanDistance(centroids[(\beta, \alpha_1)], centroids[(\beta, \alpha_2)])$
            $\delta.append(\delta_\beta)$
        **end for**
        Compute mean Euclidean distance across batch effect elements:
$D_{ref}[\alpha_1, \alpha_2] = mean(\delta)$
    **end for**
**end for**
Normalize: $D_{ref} = \frac{D_{ref}}{max(D_{ref})}$

*Continued from previous page:*

*// Step 2: Calculate Loss For Mini-Batch During Training*
Input mini-batch: *adata*
Define: $X = adata.X$ *// Model-generated embedding space from scRNA-Seq data*
Define: $A =$ "*cell_type*"
Define: $B =$ "*patientID*"
Define: $unique_A = unique(adata.obs[A])$
Define: $unique_B = unique(adata.obs[B])$
Define: $centroids = \{\}$
**for** $\alpha$ in $unique_A$ **do**
    Define mask: $mask = adata.obs[A] == \alpha$
    Apply mask and calculate centroid:
$centroids[\alpha] = mean(X[mask], axis = 0)$
**end for**
Initialize distance matrix:
$D = zeros(len(unique_A), len(unique_A))$
**for** $\alpha_1$ in $unique_A$ **do**
    **for** $\alpha_2$ in $unique_A$ **do**
        Euclidean distance between cell type clusters:
$\delta = EuclideanDistance(centroids[\alpha_1], centroids[\alpha_2])$
        $D[\alpha_1, \alpha_2] = \delta$
    **end for**
**end for**
Normalize: $D = \frac{D}{max(D)}$
Extract relevant part from reference distance matrix:
$D_{relevant\_ref} = D_{ref}[unique_A, unique_A]$
Normalize: $D_{relevant\_ref} = \frac{D_{relevant\_ref}}{max(D_{relevant\_ref})}$
$\mathcal{L} = MSE(D, D_{relevant\_ref})$ **return** $\mathcal{L}$

### 4 Attributions

Icons from flaticon.com were used to prepare some of the figures in this work.
